# Genetic diversity of maize landraces from the South-West of France

**DOI:** 10.1101/2020.08.17.253690

**Authors:** Yacine Diaw, Christine Tollon-Cordet, Alain Charcosset, Stéphane Nicolas, Delphine Madur, Joëlle Ronfort, Jacques David, Brigitte Gouesnard

## Abstract

From the 17th century until the arrival of hybrids in 1960s, maize landraces were cultivated in the South-West of France, a traditional region for maize cultivation. A set of landraces were collected in this region between the 1950s and 1980s and were then conserved *ex situ* in a germplam collection. Previous studies using molecular markers on approx. twenty landraces fo this region showed that they belonged to a Pyrenees-Galicia Flint genetic group and originated from hybridization between Caribbean and Northern Flint germplasms introduced in Europe. In this study, we assessed the structure and genetic diversity of 194 SWF maize landraces to elucidate their origin, using a 50K SNP array and a bulk DNA approach. We identified two weakly differentiated genetic groups, one in the Western part and the other in the Eastern part. We highlighted the existence of a longitudinal gradient along the SWF area that was probably maintained through the interplay between genetic drifts and restricted gene flows, rather than through differential climatic adaptation. The contact zone between the two groups observed near the Garonne valley may be the result of these evolutionnary forces. We found only few significant cases of hybridization between Caribbean and Northern Flint germplasms in the region. We also found gene flows from various maize genetic groups to SWF landraces. Thus, we assumed that SWF landraces had a multiple origin with a slightly higher influence of Tropical germplasm in the West and preponderance of Northern Flint germplasm in the East.

## Introduction

Maize was domesticated from teosinte, *Z. mays* L. subsp. *parviglumis*, about 9000 years ago in Southern Mexico (1). Thereafter, it spread from the domestication center over North and South America (1-3). During these expansions, maize evolved in contrasted environments, leading to distinct genetic groups adapted to new climates such as cold temperatures and long days in the north and warmer temperatures in coastal Central America. After the discovery of America by Columbus in 1492, maize was introduced in Europe for the first time in 1493 from the Caribbeans to Spain (4-6). In the first half of the 16th century, a second introduction of maize is documented in North-Eastern Europe from Northern Flint material originating from North America as suggested by an illustration of Leonard Fuchs, a German herbalist, in 1542 (4, 7). Genetic and morphological differences between European maize landraces were found in numerous studies. Studies based on morphological data separated European maize landraces principally according to their earliness and morphological traits also related to flowering time earliness (8, 9). Maize landraces from North-Eastern Europe were shown to be earlier than those from Southern Europe. Molecular studies confirmed the major differentiation between maize from Northern and Southern Europe (4, 7-11). Gauthier *et al*. (9) also detected, in their study, genetic differentiation between South-Western (Northern Spain, Portugal and the Pyrenees) and South-Eastern (Greece and Italy) European landraces. These results suggested that the landraces cultivated in the North, South-West and South-East of Europe resulted from introductions of different maize landraces belonging to different American regions.

Molecular studies also showed that maize landraces cultivated at intermediate latitudes in Europe originated from hybridizations between maize landraces from Caribbean, South American and North American landraces (4, 5, 7, 10-13). These authors observed genetic similarities between American Northern Flint landraces and landraces from Northern Europe, and also between Caribbean and Southern Spain landraces, showing that the primary introductions of maize landraces from America still have direct representatives in Europe. By analysing both American and European landraces with SSR markers, Camus-Kulandaivelu *et al*. (10) structured them in 7 genetic groups referred to as Mexican (MEX), Caribbean (CAR), Corn Belt Dent (CBD), Andean (AND), Northern Flint (NF), Italian Flint (ITA) and Pyrenees-Galicia Flint groups. This shows that Pyrenees and Galician landraces constitute an original landrace group compared to American genetic groups. Dubreuil *et al*. (7) found that maize landraces cultivated in Pyrenees and Galicia display no close similarity with any of the American genetic group; but based on structure analyses, these landraces were shown to be intermediate between landraces from the South of Spain and from the North of Europe. Recently Brandenburg *et al*. (12) analysed by resequencing first-cycle inbred lines directly derived from American and European landraces and showed that Northern Flint and Southern European material (notably from Spain) were the potential ancestors of Pyrenees and Galicia landraces. Nevertheless, Mir *et al*. (11) argued that landraces from Pyrenees also displayed a hybrid origin between Northern U.S Flint and Northern South-American landraces, with a predominance of the latter. Thus, all studies highlight the importance of NF origin in the climatic adaptation of Pyrenees Galicia maize. There remains questions about the geographic origin of this NF material: did it arrive directly from America or indirectly through NF landraces from the North of Europe; these two ways for introduction of NF landraces were proposed by Tenaillon and Charcosset (13) on their map of introduction of maize in Europe. Following all the above information, it would be highly beneficial to use genetic markers analyses to better understand the evolution of maize landraces located in intermediate regions of Europe. Since maize is an outcrossing species, landraces are expected to display relatively high levels of genetic diversity. To assess this diversity, molecular studies must thus include several individuals. However, for large sets of landraces, molecular characterization based on several individuals per landrace is limited by laborious and costly experimental processes. To overcome this problem, several studies on maize landraces were done on bulks of several plants per landrace, for RFLP analysis (4, 9, 14, 15) and SSR analyses (7, 10, 11, 16-19). First, Dubreuil *et al*. (14) evaluated a method based on the RFLP analysis of balanced bulks of DNA from several individuals. They found that allelic frequencies could be estimated with a high precision in bulks of 15 individuals. The major disadvantage of bulk DNA analysis was the loss of information on individual genotypes, preventing access to variation at individual level and thus to genetic parameters such as Wright’s fixation indices (*F*_IS_ and *F*_IT_). Nevertheless, bulking is highly efficient to estimate most parameter of interest regarding population genetic structure and to make inference about the evolutionary history of populations. Recently, this bulk approach was investigated on SNP markers in maize (20), leading to the development of a new method for allelic frequency estimations (21).

In this report, we focus on the South-West of France (SWF), one of the main traditional maize cultivation areas in Europe, as a case study for local scale analysis. Maize cultivation in SWF was first reported in 1626 in the ‘Béarn’ region, and in 1628 and 1637 respectively in the towns of ‘Bayonne’ and ‘Castelnaudary’ (22). Based on historical records, maize spread from its introduction in the South-West of France at the end of the 16th century and was already largely cultivated in this area (from the Pyrenees to the Garonne Valley) at the end of the 17th century (22). American hybrids were introduced from 1948 onwards in France and maize landraces were progressively replaced by commercial American x European hybrids. The origin and genetic diversity of landraces cultivated in SWF before the introduction of commercial hybrids are poorly known, although, as SWF includes the French Pyrenean region, we could expect SWF landraces to exhibit similar characteristics to that of Pyrenees Galicia landraces. The aims of this paper are (1) to assess the structure and genetic diversity of traditional maize landraces cultivated in the South-West of France until 1960-1987, and (2) to determine genetic relationships between SWF landraces with American and European maize landraces. To do so, we analysed the 50 K SNPs array diversity of 342 maize landraces, including 194 landraces collected in the South-West of France and conserved by INRA since the 1960s, as well as 148 European and American maize landraces already analysed by Camus-Kulandaivelu *et al*. (10) and Mir *et al*. (11) using SSR markers.

## Materials and methods

### Plant material

We studied one hundred and ninety-four maize landraces that had been collected between 1949 and 1987 (23, 24) in the South-West of France (SWF, S1 Table) in the two French administrative regions “Nouvelle Aquitaine” (in 5 districts) and “Occitanie Pyrénées-Mediterranée” (in 9 districts). Since their collection, these landraces have been preserved at the French Maize Biological Resource Center (CRB, Mauguio, France, https://urgi.versailles.inra.fr/siregal/siregal/grc.do) and seed lots regenerated through four successive generations of multiplication using between 100 and 200 full-sib ears for each landrace. Passport data including information about the area of collection (continent, country and district) and geographical coordinates (latitude and longitude) are available for each landrace although geographic coordinates are lacking for 36 of these 194 landraces (S1 Table).

To address questions about the origin of these landraces and to compare landraces from SWF with original American material, we used a worldwide reference panel consisting of 64 European and 73 American maize landraces (S2 Table) previously analysed using SSR markers by Camus-Kulandaivelu *et al*. (10). These authors identified 7 genetic groups for these maize landraces that they termed the Andean (AND, *n*= 12), Caribbean (CAR, *n*= 26), Mexican (MEX, *n*=21), Northern Flint (NF, *n*=39), Corn Belt Dent (CBD, *n*=17), Italian (ITA, *n*=16) and Pyrenees-Galicia (*n*=26) groups. The Pyrenees-Galicia group consists of landraces from the Pyrenees (Spain and France), from Northern Spain and from Portugal. Thus, in this study, we excluded the 22 landraces from the SWF included in those 7 groups evidenced by Camus-Kulandaivelu *et al*. (10) to avoid duplicating the SWF landraces; so we gathered the remaining landraces from the Pyrenees-Galicia genetic group, such as landraces from Portugal and Spain in a geographical landrace group named Pyrenees_Galicia_2 landrace group. To complement this reference set, we added 2 landraces from Portugal and 9 landraces from South America previously analysed by Mir *et al*. (11).

### DNA extraction and genotyping with 50K SNP array

We assessed the nucleotide diversity of the 194 maize landraces collected in South-West France using the Maize 50K SNP array developed by Ganal *et al*. (20) and bulks of DNA representing each landrace. To this aim, DNA was extracted from leaf disks collected on 15 individuals per landrace. To assess population diversity, we selected the 30,068 Panzea markers (PZE-prefix SNPs) proven suitable for diversity analyses (20) and we predicted SNPs allele frequencies using the method described in Arca *et al*. (21) briefly presented below. The approach consists in a two-steps analysis of the relative fluorescence ratio for each allele. The first step consists in determining for each SNP, whether a landrace is fixed for the A (or for the B) allele, by comparison with the distribution of ratio of A vs. B fluorescence within a set of 327 inbred lines. If the SNP is declared polymorphic within the landrace, the second step consists in estimating the allelic frequencies at this locus using a calibrated average curve established from1000 SNPs on polymorphic DNA pools controlled for their composition. Arca *et al*. (21) observed that the mean absolute error (MAE) for allelic frequencies estimation at both steps was on average 7%. For the 82 American and the 66 European maize landraces used as a worldwide reference panel, we used the SNPs database of Arca *et al*. (21) obtained using the same SNP array and following the same procedure. Combining these different datasets, we finally obtained an allele frequency database for 23,412 SNP in which diversity was assessed on 342 landraces.

### Diversity and genetic structure analysis of SWF landraces

For each landrace, we estimated the allelic richness (*A*_r_) and gene diversity (*H*_e_, (25)) using a (26) script (S. Nicolas, personal communication).

To assess the population genetic structure underlying our panel of SWF maize landraces, we used the software ADMIXTURE v1.07 (27). This method assumes the existence of a predetermined number (K) of clusters and estimates the fraction of ancestry of each accession in each of the K clusters (Q). It also infers SNP allele frequencies of the ancestral landraces (P). The software requires individual genotypic data, while our SNP analysis on DNA bulks provides allelic frequencies only. To obtain individual data, we thus simulated 5 haploid genotypes per landrace. To limit linkage disequilibrium, this analysis was performed on a subset of 2500 SNPs chosen as follows: first we divided the maize genetic map into 2500 non-overlapping windows; then, in each of these windows, we randomly selected a single SNP. We explored K values ranging from K= 1 to K=13. The likely number of genetic groups was estimated using the *DeltaK* parameter following method proposed by Evanno *et al*. (28). Each landrace was assigned to a cluster when its fraction of ancestry in this cluster was higher or equal to 0.5. For K>2, when ancestry fraction of a landrace in the different clusters was lower than 0.5, the landrace was assigned to a composite group termed “mixed”.

### Relationship between landrace genetic structure, geographic and climatic variables

To describe the genetic structure of SWF landraces, we first looked for links between the genetic structure and the geographic localization of the landraces. To this aim, we calculated pairwise genetic distances over the 194 maize landraces using the modified Rogers’s distance (29, 30). On the corresponding genetic distance matrix, we performed a principal coordinate analysis (PCoA) using R ade4 package and estimated correlations between landrace coordinates on the first axis of the PCoA and either (1) the longitude or (2) the latitude of their site of collection. This analysis was based only on those 158 SWF landraces with available geographic coordinates. We also looked for patterns of isolation by distance. To this aim, we performed a regression analysis between the “genetic divergence matrix” which we estimated using the linearized *F*_ST_ /(1-*F*_ST_) values following Rousset (31), and the pairwise geographic distance matrix calculated using latitude and longitude coordinates of the site of colection of each landraces with spherical law of cosines formula (32). Pairwise *F*_ST_ values were calculated over SWF landraces using the Gst estimate proposed by Nei (33). The statistical significance of the correlation between these two matrices was evaluated using a Mantel test ((34), R vegan package).

We also explored relationships between the genetic structure and climatic variables. To this aim, we first retrieved data for monthly precipitation and monthly minimum and maximum average temperatures from the WORLDCLIM database (35). From these, we retained 12 variables: the mean monthly temperature and mean monthly precipitation of the 6 months of maize cultivation in Europe, i.e., from May to October. We performed a principal component analysis on these 12 variables with R ade4 package. Doing this, we observed that the first axis of this PCA explained 82.5% of the climatic variability observed for temperatures and precipitations covered by our set of 158 SWF landraces. The climatic matrix distance between our 158 landraces was computed as the Euclidian distance on this first PCA axis (Ecodist R package). Thereafter, we used the function MRM of the Ecodist R package to perform a multiple regression of the genetic matrix on the geographical matrix and the climatic matrix. We also compared through an ANOVA analysis, the climatic characteristics of the different genetic groups identified using ADMIXTURE.

### Comparative analyses between SWF landraces and worldwide maize genetic groups

#### Genetic diversity

To compare allelic richness and gene diversity between well-known maize genetic groups and our set of SWF landraces, we used results obtained using ADMIXTURE on our panel of 194 maize landraces for K=2 and considered the seven genetic groups previously identified by Camus-Kulandaivelu *et al*. (10). Allelic richness and genetic diversity (*H*_e_) were estimated on each of these groups and, to compare allelic richness (*A*_r_) among groups of different sizes, we used the rarefaction method proposed by Petit *et al*. (36) with *n*=1000 resampling. To estimate the expected heterozygosity (*H*_e_) for each group we calculated the averaged values of allelic frequencies at the 23 K SNP over the landraces owing to each group. The *H*_e_ values were then averaged over SNPs. Pairwise comparisons of *A*_r_ and *He* values between groups were carried out using Wilcoxon signed-rank tests across all the 23 K SNPs. Statistical analyses were computed using a R Core Team (26) script (S. Nicolas, personal communication).

#### Looking for footprints of hybridization between main maize genetic groups in the origin of SWF landraces

To determine whether SWF landraces originated from hybridizations between ancestral landrace groups of maize s, we used the 3-populations test (37) implemented in the TreeMix software version 1.12 (38). This test compares a focus population X to two reference populations Y and W, and calculates an *f3 statistic*, f3(X; Y, W) defined as the product of the difference in allele frequencies between population X and Y, and the difference in allele frequencies between populations X and W. If the focal population X can be considered as resulting from an admixture or hybridization event between population Y and W, the value of the *f3-stat* can be negative. A *Z-score* value <-2 indicates significant mixture (37).

We also tested two scenarios for the origin of SWF landraces. In the first one, we made the hypothesis that SWF landraces originated from hybridizations between landraces of Caribbean and Northern Flint (NF) groups as proposed in previous articles. In the second scenario, we assumed that SWF landraces originated from hybridizations between maize landraces belonging to unknown groups we thus wanted to identify. For example, to determine whether SWF landraces originated from hybridization between landraces belonging from NF and MEX groups, we performed TreeMix 3-population tests using the combination of three populations analysed formed by a focal SWF landrace, with one NF and one MEX landraces as potential ancestors of the focal landraces. To analyse these two scenarios of origin, we performed all combinations of three landraces formed by (1) a focal SWF landrace (194 landraces) and two landraces from American groups; (2) a focal SWF landrace (194 landraces) and two landraces from European groups; and (3) a focal SWF landrace (194 landraces) and one landrace from an American group and one landrace from a European group. This resulted in 1,807,304 tests.

To analyse more broadly the genetic proximity between SWF landraces and worldwide maize genetic groups, we also performed a principal component analysis (PCA) with ade4 R package using SNP data from 342 maize landraces. To perform this analysis, we only used the 72 American landraces representing for the most part the 4 main historical maize genetic groups (i.e. MEX, AND, CAR and NF groups). The European maize landraces (including the SWF landraces) were added as supplementary data. Due to their recent origins, the landraces of CBD group (39) were also added as supplementary data. We used the modified Rogers’ distance between each SWF landrace and each American landrace in order to identify the closest American maize genetic group.

## Results

Among the 23412 SNPs analysed, 378 exhibited missing data for at least one landrace; they were consequently discarded from analyses leaving 23034 SNPs. Among them, 15 were monomorphic on all the landraces considered in this study (*n*=342). At landrace level, the proportion of polymorphic loci varied from 13.5% for IPK60, a German Northern Flint landrace, to 92.8% for EPZMV23, a landrace from Spain. On average, the proportion of polymorphic loci within landrace was 68.7%.

### Genetic diversity and population structure

The proportion of polymorphic SNPs in landraces from the South-West of France varied between 34% and 86%, with an average value of 70%. Averaged over SNPs, the mean allelic richness (*A*r) per landrace was 1.70, and varied from 1.33 to 1.86. Mean gene diversity (*H*_e_) was 0.22 per landrace (min = 0.12; max = 0.27, Fig 1, S1 Table).

**Fig 1.**
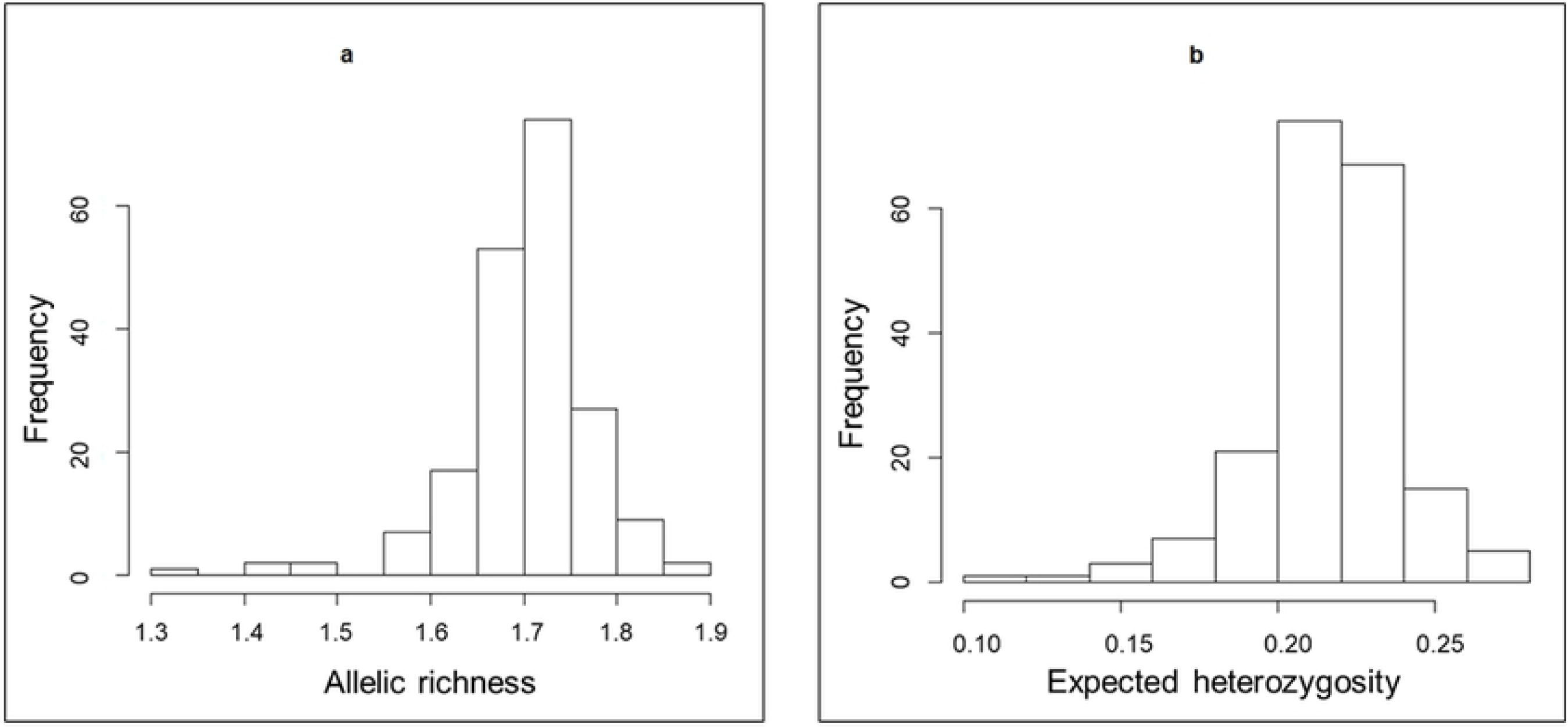
Genetic diversity analysis of SWF landraces. Histogram of (a) allelic richness and (b) expected heterozygosity values estimated for each of the 194 landraces.

Using the software ADMIXTURE on the whole set of SWF landraces, we identified a stratification in two major groups (the maximal deltaK value occurred at K=2, S1 Fig). As shown on Fig 2, the two groups distinguished landraces located in the Eastern part of the South-West of France from those located in the Western part. The first group which was named ‘East South-West France’ (S-SWF) included 65 landraces. Most of these landraces were collected in the Eastern districts such as Ariège, Tarn and Haute-Garonne; but ten of them were collected in Western districts of SWF (i.e., from districts such as Pyrénées Atlantiques, Hautes-Pyrénées, Landes and Gironde). The second group referred to hereafter as ‘West South-West France’ (W-SWF) included 126 landraces mostly originating from the Hautes-Pyrénées, Pyrénées-Atlantiques and Landes districts. Twenty-three landraces from this group were nevertheless collected in the Eastern part of the area. Interestingly, most of the landraces collected in the Haute-Garonne and Ariège districts exhibited patterns of admixture between W-SWF and E-SWF groups. At K=3, we observed a third group mostly composed of the 10 landraces that were assigned to E-SWF group at K=2 but located in the western part of the region (S2 Fig).

**Fig 2.**
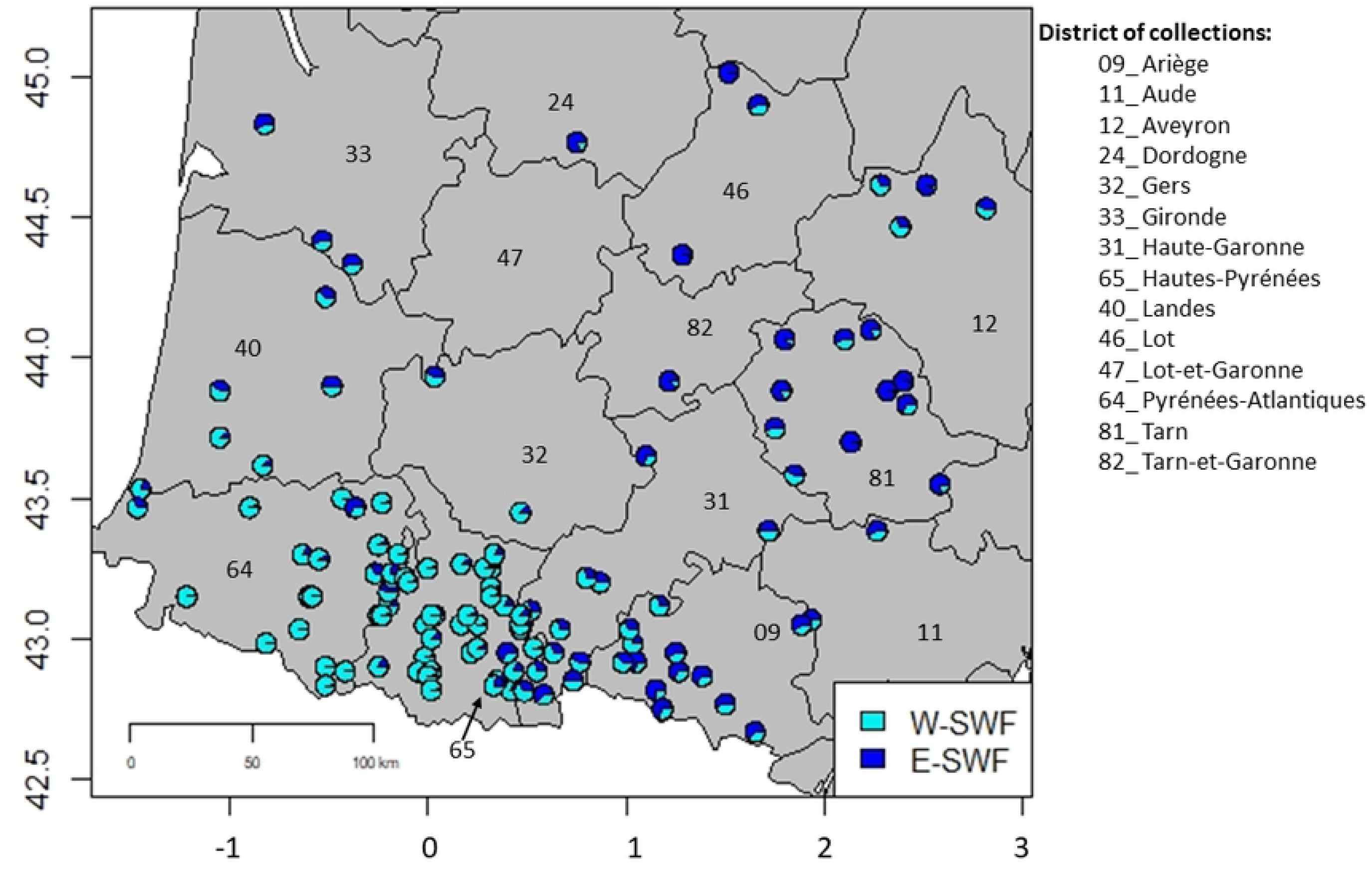
Geographical representation of the 158 maize landraces from the South-West of France (36 landraces having no geographical coordinates were not plotted). The districts of collection sites are represented with administrative numbers. Landraces for which pie-diagram had both blue and cyan colors exhibited mixed genetic origins between E-SWF and W-SWF groups.

PCoA on allele frequencies revealed the same pattern of population structure. The first axis (representing 13% of the total inertia) separated landraces located in the Western part (W-SWF cyan colour in Fig 3a) from those located in the Eastern part (E-SWF dark blue colour in Fig 3a). The second PCoA axis of Fig 3a (representing 6.3% of total inertia) highlighted variation among E-SWF landraces. Six landraces, assigned to the third group identified with ADMIXTURE at K=3 (S2 Fig) were indeed strongly differentiated from the majority of E-SWF landraces on the PCoA plan 1-2.

**Fig 3.**
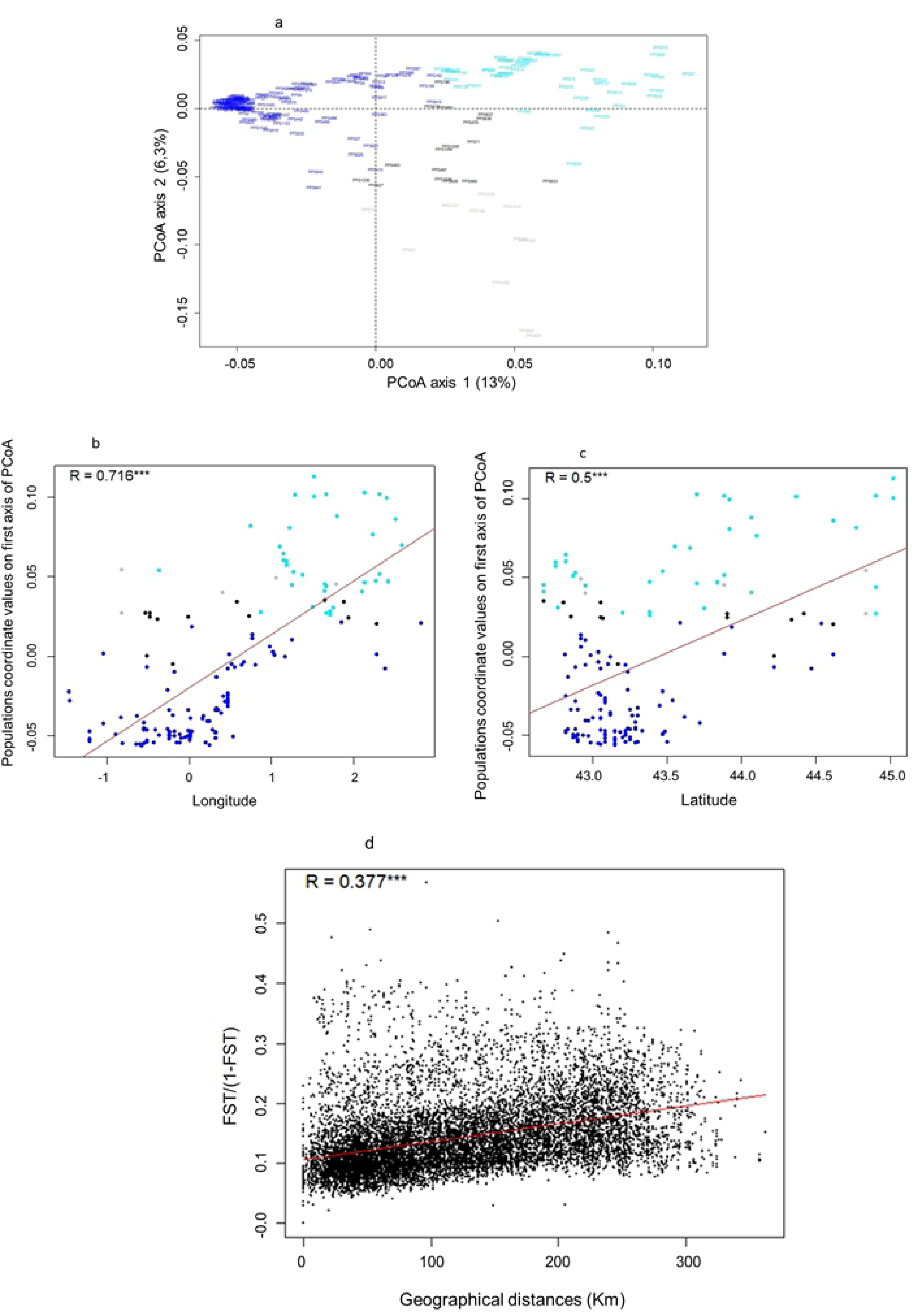
Spatial genetic structuration analysis of SWF landraces. (a)Principal coordinate analysis on SNP data using Rogers ‘genetic distance matrix estimated among the 194 landraces. (b) Longitudinal and (c) latitudinal gradient analyses on SNP data of 158 landraces from South-West France using coordinates values on the first axis of PCoA analysis. SWF landraces were colored according to their assignation to E-SWF and W-SWF groups identified with admixture analysis at K=2. (d) Isolation by distance (IBD) analysis of 158 landraces from South-West France using *F*_ST_ / (1-*F*_ST_) matrix for genetic differentiation estimated following Rousset (1997) and geographical distance matrix estimated for each pair of landraces. R= coefficient of correlation. ***= p-value<0.001.

We analysed the genetic diversity of each of these two groups. The allelic richness of W-SWF was significantly higher than that of E-SWF: *A*_r_=1.718 and *A*_r_= 1.679 respectively (Wilcoxon’s signed-rank test, *p-value* < 10^−15^; Table 1). On the other hand, the E-SWF group exhibited a higher level of gene diversity compared to the W-SWF group: *H*_e_=0.290 and *H*_e_= 0.273 respectively (*p-value* < 10^−15^ using Wilcoxon’s signed-rank test). This discrepancy between *A*_r_ and *H*_e_ indicates that allelic frequencies are more balanced in E-SWF than in W-SWF. The W-SWF/E-SWF differentiation explained a low but significant proportion of the overall genetic diversity: *F*_ST_ =0.026 (*p-value* < 10^−15^ based on permutation test over landraces). The mean *F*_ST_ between pairs of landraces was 0.104 and 0.168 for the W-SWF and E-SWF groups, respectively. This shows that the E-SWF landraces are, on average, more differentiated from each other than the W-SWF landraces.

**Table 1.**
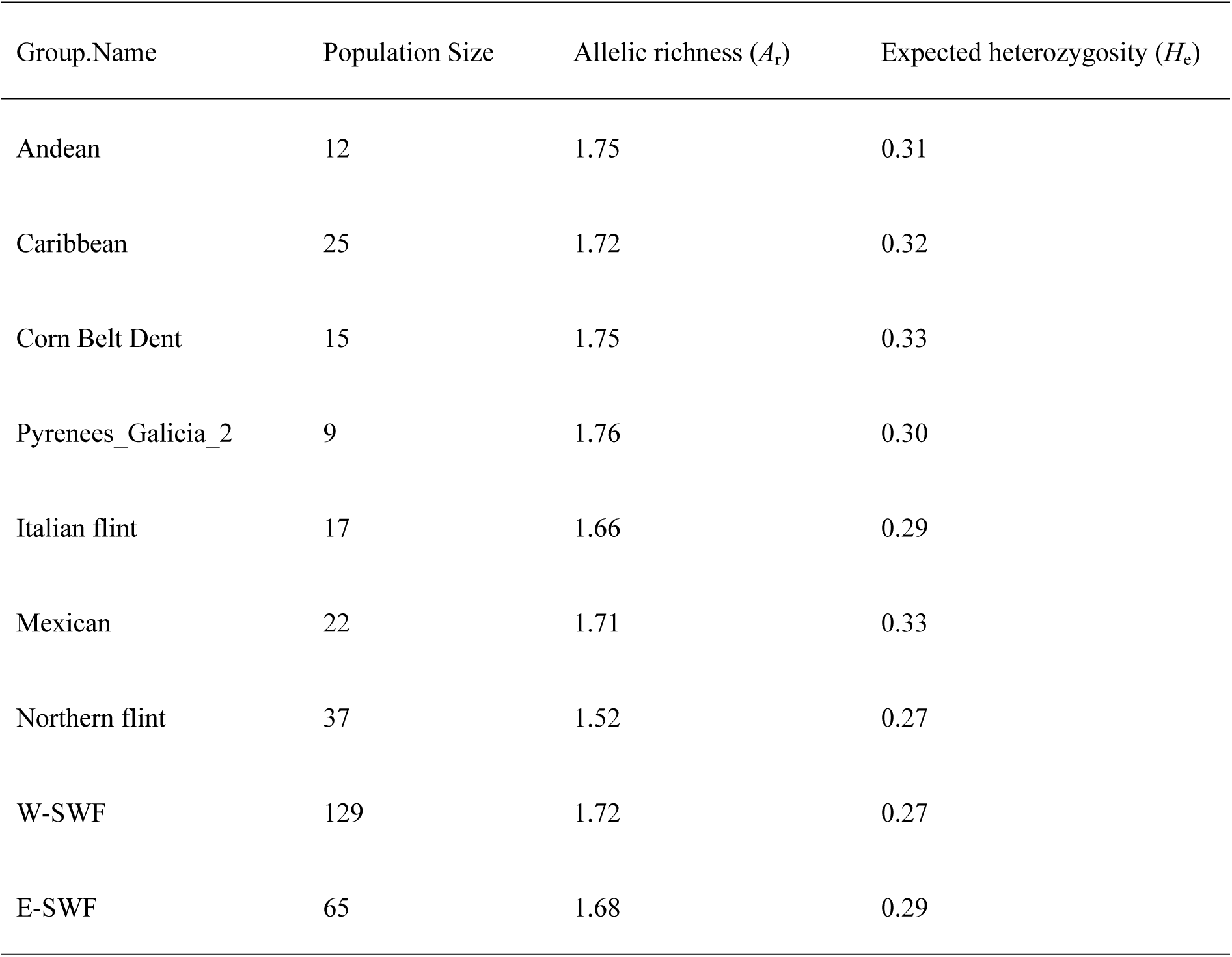
**Allelic richness (*A*_r_) and gene diversity (or expected heterozygosity, *H*_e_) for the two SWF groups identified with admixture at K=2 and the 7 genetic groups previously identified in Camus-Kulandaivelu *et al*. (10).** *A*_r_ values were calculated using rarefaction method as in (36), with 1000 resampling of 9 landraces without replacement in each group; except for Andes group for which we obtained only 220 possible resampling of 9 landraces. We used a Wilcoxon signed-rank test to analyze statistical differences between peers of *H*_e_ (or pair of *A*_r_) values estimated for the 9 groups. All comparisons for pair of *H*_e_ (and pair of *A*_r_) were significant (*p-value*< 0.009); except for comparison between He of ITA and SWEF groups (*p-value*= 0.45) and comparison between *A*_r_ of Andean and Corn Belt Dent groups. E-SWF= East South-West France; W-SWF= West South-West France.

| Group.Name | Population Size | Allelic richness ($A_r$) | Expected heterozygosity ($H_e$) |
| --- | --- | --- | --- |
| Andean | 12 | 1.75 | 0.31 |
| Caribbean | 25 | 1.72 | 0.32 |
| Corn Belt Dent | 15 | 1.75 | 0.33 |
| Pyrenees_Galicia_2 | 9 | 1.76 | 0.30 |
| Italian flint | 17 | 1.66 | 0.29 |
| Mexican | 22 | 1.71 | 0.33 |
| Northern flint | 37 | 1.52 | 0.27 |
| W-SWF | 129 | 1.72 | 0.27 |
| E-SWF | 65 | 1.68 | 0.29 |

### Relationships between geographic distribution, climate variables and population structure

To determine the main factors underlying the genetic structure observed in SWF landraces, we looked for a relationship between genetic variation and either the geographical origin of the landraces or the climatic characteristics of their site of origin. To this aim, we first looked for associations between (i) landrace coordinates on the first axis of the PCoA (Fig 3a) performed on the SNPs and (ii) either latitude or longitude coordinates of their prospection sites. As shown on Figs 3b and c, linear regression analyses evidenced significant correlations with both longitudinal (*r*=0.72; *p-value* < 10^−15^) and latitudinal coordinates (*r*=0.5; *p-value* = 10^−10^), highlighting the existence of both longitudinal and latitudinal genetic gradients. We also identified a significant correlation between the pairwise *F*_ST_ / (1-*F*_ST_) ratio and pairwise geographic distances (*r*=0.38, *p-value* < 10^−15^, Fig 3d). This result clearly marks a pattern of isolation by distance.

Examining climate characteristics from May to October in the collection sites of our set of SWF landraces, we observed that landraces belonging to the E-SWF group were from sites associated with higher temperature and weaker precipitation than W-SWF landraces (S3 Table). Differences in monthly temperatures and precipitations between E-SWF and W-SWF landraces were low, but significant varaitions were observed from June to August for temperature and during September and October for precipitations (see S3 Table, in bold). To explore how much genetic differentiation between SWF landraces could be explained by geographic distance instead of climatic distance, we performed a multiple linear regression of both geographic and climatic data matrices with the genetic data matrix. Results showed a significant relationship only for the geographic matrix (*p-value*=0.01 for geographical values VS *p-value*= 0.67 for climatic values), which suggests that the demographic history (instead of selection) of these populations is the main factor driving population differentiation.

### Genetic diversity of SWF landraces compared to the main maize genetic groups

As shown in Table1, the two genetic groups identified in the South-West of France using ADMIXTURE exhibited larger allelic richness (*A*_r_) and gene diversity (*H*_e_) than the Northern Flint group, but lower allelic richness and gene diversity than the Corn Belt Dent (CBD), Andean (AND) and Caribbean (CAR) groups. *H*_e_ values were higher for MEX than for E-SWF and W-SWF groups, whereas the *A*r of MEX was higher than the *A*r of E-SWF group and lower than that of W-SWF group.

#### Do SWF landraces originate from hybridization between known maize genetic groups?

We used treeMix 3-population test to determine whether SWF landraces result from hybridization events beween histrorical maize genetic groups. Two different scenarios of hybridization were considered. The first scenario assumes that SWF landraces originated from hybridization events between Caribbean and Northern Flint landraces. We distinguished Northern Flint landraces from North America (NFA) from those from Northern Europe (NFE) and we analysed two hypotheses: SWF landraces originate from (1) an hybridization between Caribbean landraces and Northern Flint landraces previously introduced in Europe (NFE), or (2) an hybridization between Caribbean landraces and American Northern Flint landraces (NFA) independently introduced in the South-West of France. To do this, we performed TreeMix 3-populations tests on all combinations of a focal SWF landrace with each of the Caribbean landraces included in our analysis and each of the NF landraces representing either the Northern America (NFA) or Northern Europe (NFE) groups (resulting in 43640 tests for the CAR x NFA scenario and 126100 tests for the CAR x NFE scenario; Figs 4a and b, S4 Table). Negative *f3-stat* values, suggestive of a hybridization event between CAR and NFE, were detected for triplets involving five landraces from the E-SWF group and three landraces from the W-SWF group (922 triplets. The *f3-stat* values suggestive of a hybridization event between CAR and NFA landraces were observed for the same set of SWF landraces (284 triplets). This test was thus unable to determine the implication of NFE versus NFA in hybridization events. When looking for *Z-score* < −2 (S4 Table), four landraces of the E-SWF group and PPS1236 of the W-SWF group had significant triplets with both NFA and NFE landraces; while PPS363 of the W-SWF group had significant triplets with only NFE landraces. These results suggest that only 6 SWF landraces can be considered as originating from hybridization events between landraces from NF (either Northern America or those introduced in Europe) and CAR groups The relevant CAR landraces in these analyses were located in various geographic regions: Caribbean Islands (13 landraces), Northern South America (1 landrace from Venezuela), Costa Rica (1 landrace) and Southern Spain (3 landraces).

**Fig 4.**
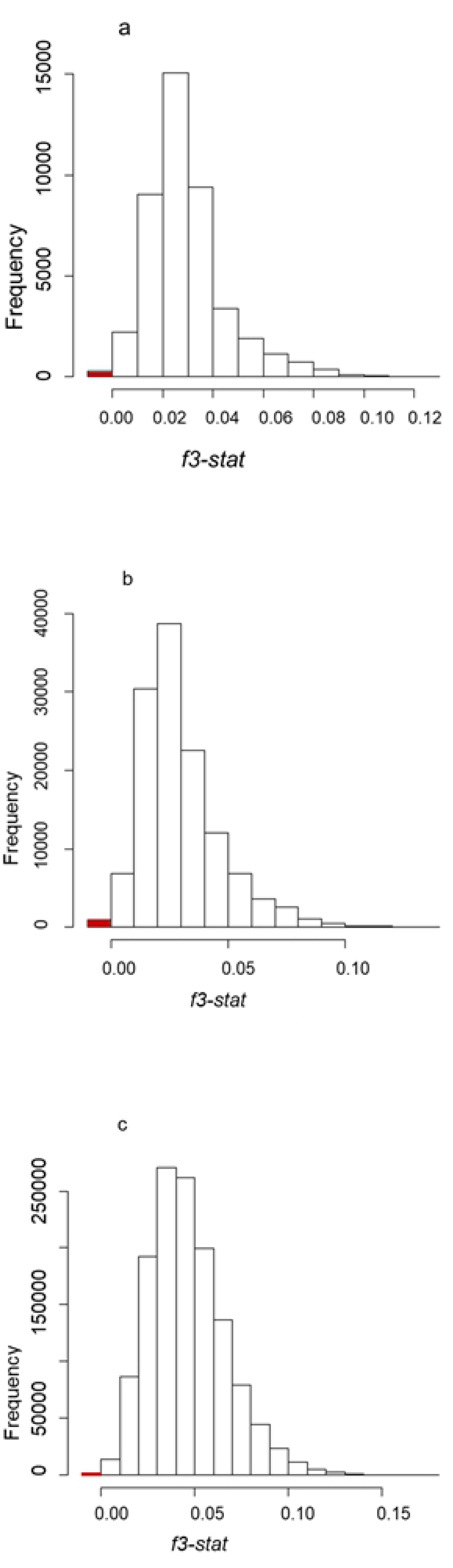
Histogram of f3-stat values estimated by TreeMix three-population test. Each f3-stat corresponds to triplet for a focal SWF landrace with two landraces from two different genetic groups such as Caribbean (CAR), Northern flint (NF), Mexican (MEX), Andean (AND), Italian (ITA), Corn Belt Dent (CBD) and Pyrenees_Galicia_2 groups. (a) Histogram for combinations of SWF landraces with CAR and American Northern flint (NFA) landraces. (b) Histogram for combinations of SWF landraces with CAR and European Northern flint (NFE) landraces. (c) Histogram for other combinations. Red color indicated negative values for triplets of landraces, showing that some SWF landraces could have originated from mixture events of landraces from other genetic groups.

In a second step, we used three-mix texts to test whether SWF landraces originated from hybridizations between any of the historical genetic groups. This led us to analyse 1329094 different triplets. Over those, only 204 exhibited both negative values for the *f3-stat* parameter and *Z-score*<-2 (Fig 4c, S4 Table). Interestingly, all these triplets included one NF landrace as one of the ancestor of the SWF landraces (S4 Table), the second ancestral landrace belonging either to the MEX, AND or ITA groups. For SWF landraces, we detected such a hybridization footprint for only 4 landraces. These 4 landraces from the South-West of France were also implied in the triplets showing signal of hybridization events between CAR and NF landraces (S4 Table). We also observed 4 other SWF landraces that showed signals for hybridization events between landraces from at least two genetic groups. It concerned (1) 2 landraces of E-SWF showing signals for hybridization events between landraces from only NF and AND groups (PPS1248 and PPS1249); (2) one landrace of W-SWF showing signals for hybridization events between landraces only from NF and MEX groups; and (3) 1 landrace of W-SWF (PPS616) showed signals for hybridization events between landraces of NF and landraces from either MEX or ITA groups (S4 Table). All these results suggested that few landraces from the SWF (1) originated from hybridization events between landraces from NF and either CAR or MEX or AND maize genetic groups, (2) the NF germplasm played important role in the evolution of these SWF landraces and (3) maize landraces of ITA group showed genetic similarity with some SWF landraces.

We did not observe signals for hybridization events including CBD landraces as one of the ancestors of the SWF landraces, suggesting either that these were not influenced by CBD landraces after their introduction in Europe, or that we could not evidence it with the TreeMix 3-population test.

#### Genetic proximity between SWF landraces and maize from Europe and America

As we detected only few clear evidences of hybridization events in SWF landrace origins using a TreeMix 3-population test, we performed a PCA on SNPs for the whole set of 342 maize landraces (Fig 5, S2 Table). Fig 5 shows the position of the different groups on the first and second axes of the PCA. The first axis (19.7% inertia) mainly separated Northern Flint landraces from Tropical landraces (AND, CAR, MEX) while the second axis (6.4% inertia) mainly separated South American from Caribbean landraces. As expected, we observed that most European NF landraces were close to American NF landraces, and that landraces from Southern Spain were close to those from the Caribbean Islands. The two SWF groups occupied a central position on the first PCA plan (Fig 5) but somewhat separated on the two first axes. Landraces of the E-SWF group were more dispersed compared to W-SWF landraces that appeared more grouped. On the first axis, E-SWF landraces appeared closer to the NF group than did W-SWF landraces. On the contrary, W-SWF landraces were closer to tropical landraces (AND, CAR and MEX) than to NF landraces (Fig 5). Looking for relationships between landraces from Europe, we found that landraces from the SWF were closer to Pyrenees_Galicia_2 and European Northern Flint landraces (NFE) than to Italian flint landraces.

**Fig 5.**
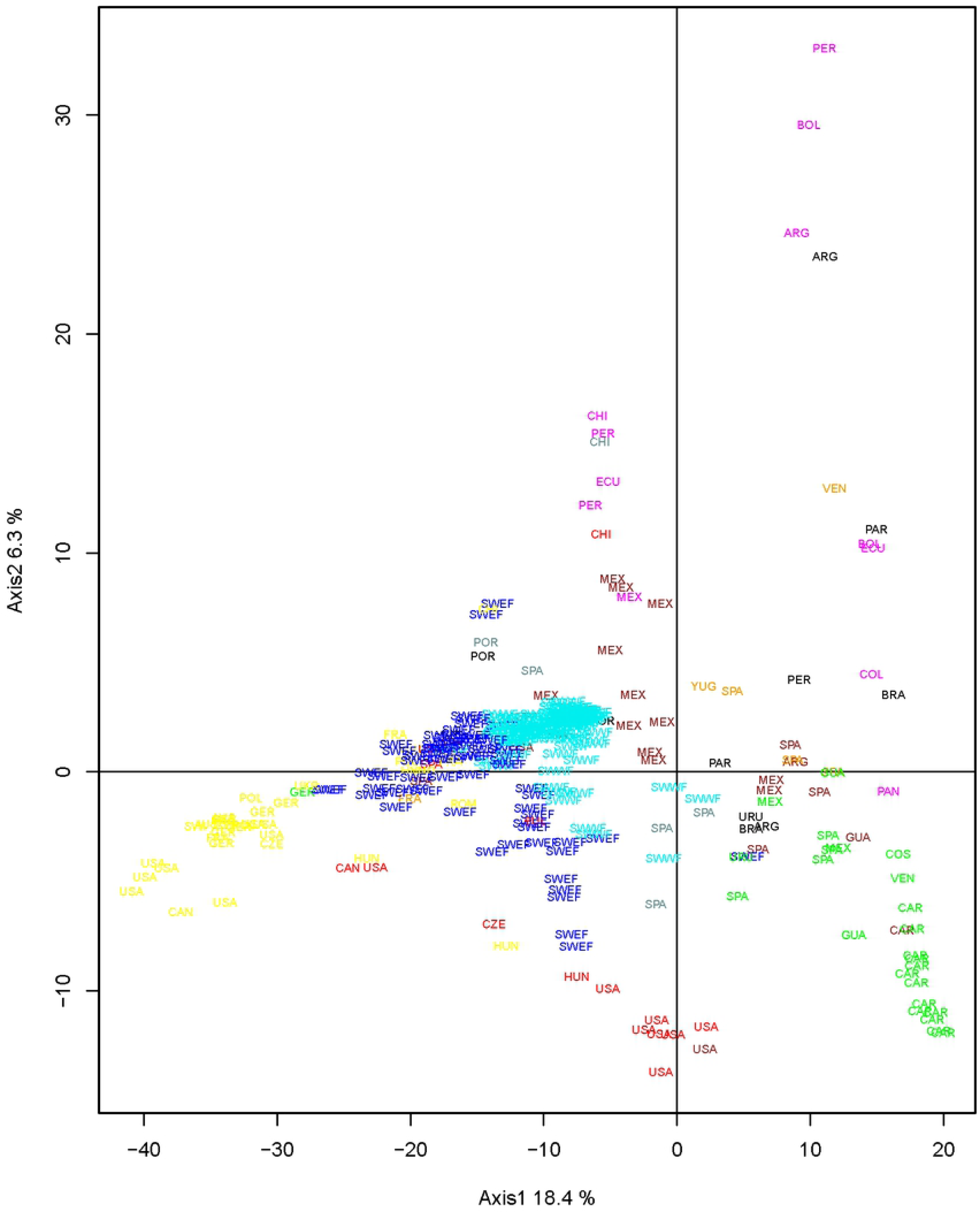
PCA analysis on SNP data of 72 American maize landraces with 260 maize landraces from Europe and those of CBD as supplementary data. Labels indicate landrace country of origins (see Table_S2). South-West France landraces were colored according to their admixture result at K=2 allowing to distinguish East (in blue) and West South-West France (in cyan) genetic groups. The remaining American and European landraces were colored according to their genetic groups previously identified by Camus-Kulandaivelu et al. (2006): Corn Belt Dent in red, Caribbean in green, Northern flint in yellow, Mexican in brown, Italian Flint in orange, Andean in magenta, Pyrenees_Galicia_2 in grey. The 9 landraces from South America and 2 landraces from Portugal studied by Mir et al (2017) were colored in black.

As a second way to identify genetic proximity between landraces, we used a heatmap representation with R gg2plot function to visualize the result of genetic distance estimation between SWF landraces and the 82 American landraces (S3 Fig). The heatmap representation of genetic distances showed that the 194 landraces from SWF were closer to 4 Chilean (PPS949 from AND, PPS941 from CBD, PPS961 from NF and PPS938 from MEX groups), 2 CBD (PI214189 and PI280061) and 2 NF (PI213793 and PI401755) landraces from North of America than landraces from others American countries. These results suggested an introduction of Northern Flint and CBD landraces from Northern America into the SWF genepool, as well as exchanges of maize landraces between Chile and SWF. Comparing the genetic distances of the two SWF groups with the American accessions, we did not evidence any clear difference between the genetic origins of the two groups from the South-West of France, except that E-SWF landraces appeared to be close to landraces from the NF group and that W-SWF landraces were closer to Tropical landraces (S3 Fig).

## Discussion

### Validity and limitations of bulk DNA analysis with the 50 K SNP array

The usefulness of the bulk DNA approach in genetic diversity investigations has been proven in numerous studies (8, 14, 17, 18, 21, 40). The major constraint observed for the use of this pooled DNA analysis is the loss of information about genetic variation at the individual level (8, 15). As a result, genetic parameters such as individual’s heterozygosity and Wright’s F_IS_ fixation indices (30, 33) cannot be estimated directly. Bulk analysis also limits the use of classic genetics tools that require individual information such as the softwares ADMIXTURE (27), GENEPOP (41), HZAR (42) and STRUCTURE (43). As a result, we chose to estimate genetic parameters and to use software tools that do not require individual genotype information to characterize and analyse these landraces, except for the use of the ADMIXTURE software for which we simulated genetic haplotypes (27).

Despite these constraints, the genetic proximities we revealed between American and European landraces using the 50K SNPs on DNA bulks were in accordance with what is known about the main paths of maize introductions in Europe. Indeed, our PCA analysis allowed distinguishing the main historical maize genetic groups previously identified by Camus-Kulandaivelu *et al*. (10). For instance, the genetic similarity between maize landraces from the South of Spain and Caribbean landraces corroborates the introduction of maize by Columbus from the Caribbean Islands to Spain (4-6). We also detected strong genetic proximity between Northern Flint and landraces from the North of Europe, in accordance with the introduction of maize landraces from Northern America to Northern Europe as suggested by an illustration of Leonard Fuchs, a German herbalist, in 1542 (4, 7).

### Diversity and genetic structure of maize landraces from the SWF

Using a genetic database of 23412 SNP markers to analyse the genetic diversity of maize landraces that were collected in the South-West of France circa 50 years ago, we were able to identify two main groups of landraces, one located in the Eastern side and one in the Western side of the South-West of France. However, although these two groups were identified through different analyses (ADMIXTURE, PCoA), the genetic differentiation between them was relatively low (*F*_ST_ = 0.03) and indicated that each group contains an important genetic variability.

The geographical repartition of the two groups and their low divergence led us to propose two hypotheses for the origin and evolution of maize landraces in the South-West of France. The first hypothesis was that these landraces originated from the same ancestral group of landraces and were then differentiated into 2 groups under the effect of genetic drift and reduced gene flow. This hypothesis was supported by the results obtained in our analysis of the pairwise genetic distance matrix between landraces, which revealed patterns of isolation by distance (IBD). We highlighted such a longitudinal gradient along SWF using a multivariate technique (PCoA) based on a genetic distance matrix with SNP data from 158 landraces, which led to a significant correlation coefficient between the first PCoA axis and the longitudinal coordinates of their prospection sites. Multivariate approaches have been used in many studies to analyze environmental or genetic gradients in landraces analysis (44-46). For their study of human genetic variation, Menozzi *et al*. (45) presented a synthetic map with the use of principal components (PCA) to condense information from many alleles and concluded that genetic gradients were suggestive of historical migrations across continents. However, Novembre and Stephens (46) found that PCA correlating with geographical did not necessarily reflect specific migration events but may instead have simply reflected isolation by distance.

The second hypothesis was that each of the identified genetic groups originated from a distinct ancestral group of maize landraces that were introduced in SWF. This second hypothesis was however invalidated by the low level of differentiation observed beween the two groups. Our results therefore support the conclusions of a previous analysis on some of these populations using SSR data suggesting that SWF landraces originated from a single genetic group of maize closest to the ancestors of the Pyrenees-Galicia group (*F*_ST_ =0.03 between the Pyrenees_Galicia_2 group and each of the goups from the South-West of France) (4, 7, 8, 10).

The geographical distribution of maize landraces in SWF could also be the result of a diversifying selection. The occurrence of a genetical longitudinal gradient along the Pyrenees area as observed in this study has been reported for several species (47-50). Gradients, such as altitude or longitude, play an important role in the Pyrenees, as they shape the existing proportion of suitable habitats along the mountains (48, 49). Valbuena-Ureña *et al*. (50) emphasized that longitude is an important gradient to take into account in the Pyrenees, because the influence of the Atlantic ocean provides cooler and wetter climate westwards than in Eastern areas, which are more influenced by the Mediterranean climate.

We found differences for temperature and precipitation between Eastern and Western parts of the SWF, but regression analysis showed that climatic variations did not explain genetic variability observed between SWF landraces, when considering geographical covariates. Since climate variation are geographically structured with longitude and latitude, we still cannot exclude completely the involvement of climatic variation in the differentiation of SWF landraces into two genetic groups; our results rather suggest that demography (restriction of gene flow and genetic drift) is the main driver of population differentiation between maize landraces in this region. This is supported by the longitudinal and latitudinal gradients that we observed on the first axis of PCA, which is determined by a large number of SNPs. However, the genetic differences observed for SWF landraces could be also explained by impact of farmer’s practices. The management of seeds and farmer’s exchanges in different valleys limited by hilly mountains probably contributed to differentiate SWF landraces in different districts. Already, the ethnobotanical survey realized in 2012-2013 with farmers in the Pyrenean area did not reveal major differences in farmer’s practices between the East and West parts of the Pyrenees (51).

### The use of genome wide analysis to infer the origin of SWF landraces

#### The role of hybridization

Building on previous works, our first assumption was that SWF landraces originated from a hybridization between maize landraces belonging to Caribbean and Northern Flint genetic groups and that the hybridization occurred earlier in Spain before introduction of maize in the South-West of France. Several studies on genetic diversity of European and American maize landraces (Dubreuil et al., 2006; Rebourg et al., 2003; Revilla Temiño et al., 2003) highlighted a Pyrenees-Galicia flint group made up of European landraces including a small sample of landraces from the South-West of France and landraces from the North of Spain. This original landrace group was hypothesized to originate from hybridizations between landraces of Caribbean and Northern flint groups; however less than 30 landraces from Pyrenees-Galicia were used in these previous studies. In our study, using a much larger data set of 194 maize landraces from the SWF, the origin for SWF landraces was more precisely analysed. Our PCA analysis showed genetic proximity between landraces from the Western part of the South-West of France and some maize landraces from Spain and Portugal, supporting genetic relationship mentioned between landraces from Pyrenees and Galicia in previous studies of maize landraces named as Pyrenees-Galicia flint group (Camus-Kulandaivelu et al., 2006; Dubreuil et al., 2006; Gauthier et al., 2002). Based on historical documents, the introduction of maize in the South-West of France was reported by Renoux (22) as originating from Spain. In her study, Vouette (52) reported the name of maize in the Pyrenees valleys as “wheat from Spain”, as given by Bonnafous (1836). Similarly, the first mention of maize in the mercuriales of Verfeuil (near Toulouse) in 1637 indicates the name of “millet of Spain” (53). The vernacular names do not allow finding the genetic origin of the maize landraces, but gave indications of where the maize landraces came from before their introduction in a given country. The introduction of SWF maize from Spain in the 17th century is also justified by the coasting trade practiced by fisherman from fishing ports of Altanlic coast of France and Spain (Mauro, 1968). In his work on relations between Spain and the South of France in the 17th century, Mauro (54) cited the route of maize that left from the West part to progress towards the East part of South-West of France. The progression of maize from West to East is consistent with the greater genetic allelic richness observed in the W-SWF group than in the E-SWF group. The lower allelic richness of E-SWF compared to the former group can also be explained by their proximity to NF which is the less diverse genetic group as observed in our allelic richness and gene diversity analysis and as found by Gauthier *et al*. (9) and Dubreuil *et al*. (7).

#### The influence of Northern Flint landraces

NF landraces played an important role in the evolution of SWF maize landraces and more generally in the adaptation of maize in Europe (4, 6) and in the development of CBD germplasm (55). Our TreeMix 3-population analysis allowed us to confirm Northern flint landraces influence in the evolution of SWFlandraces. For all admixture events highlighted in this analysis, SWF landraces displayed a mixed origin between NF landraces with either CAR, AND, MEX or ITA landrace groups. There are two possible ways to explain the presence of NF marks in the genome of SWF landraces: either directly from a secondary introduction from North America (NFA), or introduced from the North of Europe (NFE) as reviewed by Tenaillon and Charcosset (2011). Revilla et al (1998) supported the NFA hypothesis by suggesting that NF landraces could have been introduced into the North of Spain at several times from the beginning of the 17th century. Later, Brandenburg et al. (2017) found that European flint inbred lines, i.e. first-cycle inbred lines directly derived from landraces after few generations of selfing, were issued from an admixture between inbred lines belonging to European Northern Flint landraces and those derived from Southern Spain lines. Our TreeMix 3-population test was not able to distinguish the involvement of NFA versus NFE in the origin of SWF landraces. This is probably due to the genetic similarity between NFA and NFE (*F*_ST_ =0.04). However, our PCA analysis showed a closer genetic proximity between maize landraces from the Eastern part of SWF and Northern flint landraces located in the North of Europe (NFE), supporting the scenario involving some gene flow from European Northern flint landraces.

#### The influence of maize landraces from the south of America

We found genetic relationships between maize landraces from SWF and maize landraces from the South America. Mir *et al*. (11) showed that maize landraces from SWF had a Northern South America origin, but our study does not allow us to really assess this path of introduction, as only four maize landraces from the North of South America were used in this studies: VEN405 from CAR, VEN736 from ITA, ANTI392 and PANA168 both from AND groups. These landraces were also found to be probable ancestors for SWF landraces in TreeMix 3-population test (Supp. Table 4); but our PCA analysis showed only evidence for genetic similarity between W-SWF landraces and those from the Northern South-America. It is possible that our study did not capture very well genetic relationships between SWF landraces and these expected Northern South-American parental landraces identified by Mir *et al*. (11), because of a limited sample of maize landraces from South America. In consequence, we had difficulty to evidence relationships between SWF landraces with Northern South America landraces. Note that American landraces introduced in Europe evolved in America during about 500 years, which limits the ability of TreeMix 3-population test to detect hybridization events between SWF and Northern South American landraces.

Our collection of landraces from SWF presented the lowest genetic differentiation with landraces from Chile (except for CHZMO8050) compared to the remaining landraces from America. All these Chilean landraces have been shown to originate from different genetic groups (10), with a probable replacement of traditional landraces with relatively recent introduction of Northern US materials (3). Genetic relatedness between Chilean and SWF landraces may be due to the introduction of SWF genepools into Chile or vice-versa. Interestingly, the Spanish (especially Basques and Andalusians) and also other Europeans such as French people (mainly coming from SWF) immigrated mainly to Chile in the second half of the 19th century, although Basque presence in Chile began in the conquistador period. Also exchanges between the Basque country and the South West of France have taken place in both directions and for a few centuries (https://en.wikipedia.org/wiki/Immigration_to_Chile, march 2020).

#### The influence of CBD landraces

Finally, our PCA analysis showed genetic proximity between 10 landraces from SWF and landraces belonging to the Corn Belt Dent genetic group (CBD). CBD is a group of maize landraces identified in previous studies as resulting from hybridizations between Northern Flint and U.S Southern Dent materials (39, 55). The presence of CBD genomic traces in SWF can be explained by the recent introduction of American hybrid seeds in Europe, since hybrids are CBD. Indeed, from 1948, with the Marshall plan, the first American hybrids were tested in France (22). Then, from 1957 onwards, the first hybrids created by INRA were developed in France; they were double hybrids with at least one American line. Farmers ‘adoption of hybrids was fairly rapid (56). These 10 landraces, showing a genetic proximity with CBD group on the PCA analysis, shaped a specific group with admixture analysis at k=3. Introgression of CBD germplasm into local landraces has also been observed in Central Italy with very variable effects (57).

#### The different scenarios for SWF landrace origin

More generally, it is thought that the South-West of France has been submitted to several introductions of maize of various genetic origins, as already reported for Spanish maize landraces (6) and in Portugal (11, 58). Evidence for subsequent admixture between the genetic groups involved in these introductions were observed in this study with TreeMix 3-populations tests, PCA analyses and genetic distance analyses. In our TreeMix 3-population test, we found only few landraces from the South-West of France that originated from hybridization events between American and European landraces. Signals for hybridization events occurring in SWF landraces could probably have been obscured by the combined effect of the high genetic diversity withing landraces and the relatively low *F*_ST_ between landrace genetic groups. The fact that SWF landraces were already hybridized before their introduction in the South-West of France could have also influenced TreeMix analysis results, as we initially had assumed that SWF landraces were similar to the landraces of the Pyrenees-Galicia group.

All these results above allowed us to describe at least two scenarios of origin that take into account the influence of Northern Flint landraces from the North of Europe predominently in the Eastern part and the genetic proximity of W-SWF landraces with Tropical landraces. For the first scenario, we postulated that, after the introduction of hybridized maize landraces from Spain (Caribbean x Northern Flint landraces), the NFE landraces spread from the North to the Eastern part of SWF where they hybridized together. The arrival of NFE landraces could have been followed by successive gene flows of maize landraces from Mexican and Andean groups introduced from America, through the Atlantic harbours to the west part of SWF.

In the second scenario for the origin of SWF landraces, we hypothesized that the first maize landraces introduced in the South-West of France displayed predominantly Northern Flint ancestries as do Galician landraces (11). Thus, these landraces spread from the Western to the Eastern parts of the South-West of France. Thereafter, Tropical landraces (from Mexican, Caribbean and Andean genetic groups) were introduced in the Western part of SWF. Presence of genetic marks from Tropical landraces also occurred through seed exchanges between Mediterranean trademen from Italy, France, Spain and Portugal at the end of the sixteenth century (11, 12). Revilla Temiño *et al*. (5), Patto *et al*. (58), Brandenburg *et al*. (12) and Mir *et al*. (11) found that the South of Europe has been submitted to maize landraces introductions from both the Caribbean islands and South America.

## Conclusion

Assessing the population genetic structure and diversity of a collection of crop accessions is the best way to develop an efficient management strategy and to improve breeding programs. Here we used DNA bulk analyses with a 50K SNP array to investigate the diversity and population structure for a panel of 194 maize landraces collected 30 to 60 years ago in the South-West of France. In this study, we assumed that the maize landraces from the South-West of France originated mostly from hybridizations between landraces of Caribbean and Northern flint groups. Introduced around 1626 through Atlantic harbours such as Bayonne, these SWF landraces have probably been submitted posteriously to gene flows from landraces belonging to various countries of America and Europe, as expected by Revilla Temiño *et al*. (5) for maize landraces from Spain and Patto *et al*. (58) for those from Portugal. A single mixture of NF and CAR landraces is not sufficient to explain the genetic origin of SWF maize.

The differentiation between W-SWF and E-SWF (*F*_ST_=0.026) observed among SWF landraces with admixture analysis can result from demographic processes with influences of gene flow (isolation by distance) and genetic drift. Adaptation processes related to farmer’s practices could not be completely excluded. This would require a study associating genetic studies and morphological characterization to ethnobotanical surveys with old farmers who still remember maize cultivation in the 60s.

## Acknowledgements

We are grateful to Dr Ndiaga Cissé (CERAAS/ISRA, Thies, Senegal) for providing financial support, all colleagues of GE2POP staff (UMR AGAP-INRAE, France) for reviewings and orientation, Anne Zanetto (French maize Biological Ressources Center, Montpellier, France) for providing landraces. This work was supported by the CIRAD - UMR AGAP HPC Data Center of the South Green Bioinformatics platform (http://www.southgreen.fr/).

## List of supplementary figures

**S1 Fig. Plot of DeltaK of the log likelihood from the admixture analysis on SWF landraces SNP data.** Group numbers varied from K=1 to K=13.

**S2 Fig. Bar-plot for K=2 to K=5 for admixture analysis with the 194 SWF maize landraces.** At K=2, we differentiated W-SWF (in cyan) and E-SWF (in blue) genetic groups. At K=3, 15 landraces (in dark-grey) previously assigned to E-SWF group at K=2 constituted a group distinguished from E-SWF and W-SWF groups. At K=4, we observed a fourth group consisting of about 10 landraces located principally in “Lot” and in “Lot et Garonne” districts. At K=5, landraces from Gironde (in turquoise) differ from W-SWF groups; Landraces from “Lot” and “Lot et Garonne” districts were integrated again in E-SWF group and we observed differentiation between landraces from the third group at K=3.

**S3 Fig. Heat-map of Rogers ‘genetic distance values between the 194 landraces from SWF and the 82 landraces from America.** American landraces were sorted by latitude of their collection sites and colored as per their genetic groups previously identified by Camus-Kulandaivelu et al. (2006). We also sorted SWF landraces using their ancestries values on W-SWF group obtained with admixture analysis at K=2; thus SWF landrace numbers from 0 to 65 represent the E-SWF landrace group and SWF landrace numbers from 66 to 194 represent W-SWF landrace group. Corn Belt Dent in red, Caribbean in green, Northern flint in yellow, Mexican in brown, Italian Flint in orange, Andean in magenta and the 9 landraces from South America studied by Mir et al (2017) in black.

## List of supplementary tables

**S1 Table. Combination of passport data (column 1 to 5), genetic diversity analysis (column 6 and 7) and admixture analysis at K=2 (column 8 to 10) performing on SNP data of the 194 maize landraces from the South-West of France.** W-SWF= “West South-West France»; E-SWF =“East South-West France”.

**S2 Table. Passport information and PCA analysis result on SNP data of 342 maize landraces.**

**S3 Table. Climatic variations between E-SWF and W-SWF groups from May to October**. This table contained mean of temperature and mean of precipitation for W-SWF and E-SWF groups from May to October. We shaded in bold months for which differencies for mean temperatures between the two groups were significant (*p-value*<5%). Same thing was doing for precipitation.

**S4 Table. TreeMix 3-population test results**. This table represents a summary of TreeMix 3-population test results performed using all combinations of three landraces shaped by one landrace from SWF considered as tested population, and two other landraces (pop2 and pop3) considered as potential ancestors of the tested landraces. We only represented the results of three landraces for which *f3-stat* values were negative with *Zscore<-2*. E-SWF= East South-West France, W-SWF= West South-West France.

